# High growing season temperatures limit winter recovery of grapevines from *Xylella fastidiosa* infection – implications for epidemiology in hot climates

**DOI:** 10.1101/2022.11.02.514902

**Authors:** Lindsey Burbank, Mark S. Sisterson, Wei Wei, Brandon Ortega, Nathaniel Luna, Rachel Naegele

## Abstract

Management of widespread plant pathogens is challenging as climatic differences among crop growing regions may alter key aspects of pathogen spread and disease severity. *Xylella fastidiosa* is a xylem-limited bacterial pathogen that is transmitted by xylem sap feeding insects. Geographic distribution of *X. fastidiosa* is limited by winter climate and vines infected with *X. fastidiosa* can recover from infection when held at cold temperatures. California has a long history of research on Pierce’s disease, and significant geographic and climatic diversity among grape-growing regions. This background in combination with experimental disease studies under controlled temperature conditions can inform risk assessment for *X. fastidiosa* spread and epidemic severity across different regions and under changing climate conditions. California’s grape growing regions have considerable differences in summer and winter climate. In northern and coastal regions, summers are mild and winters cool, conditions favoring winter recovery of infected vines. In contrast, in inland and southern areas summers are hot and winters mild, reducing likelihood of winter recovery. Here, winter recovery of three table grape cultivars (Flame, Scarlet Royal, and Thompson seedless) and three wine grape cultivars (Sauvignon Blanc, Cabernet Sauvignon, and Zinfandel) were evaluated under temperature conditions representative of the San Joaquin Valley, an area with hot summers and mild winters that has been severely impacted by Pierce’s disease, and contains a large portion of California grape production. Mechanically inoculated vines were held in the greenhouse under one of three warming treatments to represent different seasonal inoculation dates prior to being moved into a cold chamber. Winter recovery under all treatments was generally limited, but with some cultivar variation. Given hot summer temperatures of many grape-growing regions worldwide, as well as increasing global temperatures overall, winter recovery of grapevines should not be considered a key factor affecting *X. fastidiosa* spread and epidemic severity in the majority of cases.

## Introduction

A major challenge for the cultivation of long-lived perennial vine and tree crops is spread of economically damaging insect-transmitted plant pathogens (Cambra et al. 2006; Gottwald 2010; Krugner et al. 2019). For plant pathogens that are geographically widespread, management approaches must be tailored to characteristics of the local environment (Gruber and Daugherty 2013). For example, vector species composition and distribution of crop and non-crop habitats that serve as sources of vectors and/or inoculum may vary with geographic area (Cornara et al. 2019; Krugner et al. 2019). Similarly, climatic differences among crop growing regions, and varied disease susceptibility among crop species and cultivars may alter characteristics of pathogen spread (Daugherty et al. 2017; Rathé et al. 2012). Ideally, disease management strategies should be based on results of replicated field trials representative of the regions and crop varieties most impacted. Because field trials are costly and time consuming, field trials are often conducted across a limited portion of the pathogens range, and with limited cultivar diversity. Accordingly, tailoring management approaches to local conditions requires predicting how differences in climate or cultivar traits may affect pathogen spread, often based on incomplete information (Deyett et al. 2019; Gruber and Daugherty 2013; Daugherty et al. 2017). Additionally, even when field studies have been conducted in a specific area, climate change may impact relevance of the results once a significant amount of time has passed.

The bacterial plant pathogen *Xylella fastidosa* causes disease in several vine and tree crops including grape, almond, peach, plum, citrus, and olive (Hopkins and Purcell 2002; Sicard et al. 2018). Depending on geographic region and crop, losses from *X. fastidiosa* can be minimal or devastating (Perring et al. 2001; Sisterson et al. 2012, 2020; Krugner et al. 2014; Saponari et al. 2019). The pathogen is transmitted by several species of xylem sap consuming insects, with key vector species depending on geographic location (Cornara et al. 2019; Krugner et al. 2019). Geographic distribution of *X. fastidious* is also limited by climate (Purcell 1980), with climate models typically including parameters regarding summer and winter temperatures as predictors of range, as well as pathogen subspecies (Hoddle 2004; Godefroid et al. 2019; Giménez-Romero et al. 2022; Godefroid et al. 2022). However, within the known range of *X. fastidiosa*, there are also differences in climate, vector dynamics, and management practices that can significantly impact epidemiology. This study focuses on *X. fastidiosa* subspecies *fastidiosa* in grapevine (Pierce’s disease) in California, and the broader insights that can be gained from combining epidemiological and climate information with experimental disease studies under controlled temperature conditions.

Grapevines exposed to cold temperatures can recover from *X. fastidiosa* infection, an observation that initially highlighted the role of environmental conditions in Pierce’s disease (Purcell, A.H. 1977). Several field and laboratory studies have investigated recovery of *X. fastidiosa*-infected grapevines exposed to winter climates found in California (Lieth et al. 2011; Feil and Purcell 2001; Feil et al. 2003). Similar studies have investigated winter recovery of *X. fastidiosa* infections in almond (Cao et al. 2011; Ledbetter et al. 2009). While there is broad support for the phenomenon of winter recovery of grapevines infected with *X. fastidiosa*, studies of this aspect of Pierce’s disease epidemiology have only been conducted with a small number of locations and grapevine cultivars, limiting the ability to make broader predictions across other regions.

Three factors are broadly shown to affect winter recovery of grapevines infected with *X. fastidiosa:* inoculation date, location (i.e., climate), and cultivar (Table 1). Purcell (Purcell 1981) and Feil et al. (Feil et al. 2003) demonstrated that there was an effect of inoculation date on winter recovery, with vines inoculated early in the season less likely to recover than vines inoculated late in the season. Thus, vines inoculated earlier in the season are presumed to have quantifiably different *X. fastidiosa* infection status (bacterial population and/or distribution throughout the plant) at dormancy than vines inoculated later in the season. In fact, monitoring of vines chronically infected with *X. fastidiosa* found that detection of *X. fastidiosa* increased as summer progressed and subsequently decreased as vines entered dormancy (Sisterson et al. 2020), indicating that *X. fastidiosa* populations rise and fall through the season. In this study, three inoculation treatments are compared: June inoculation with 8-week warming period, August inoculation with 8-week warming period, and 16-week warming period. Warming period refers to the time between inoculation and start of cold temperature exposure during which plants accumulate degree days dependent on the ambient temperature. These treatments are designed to evaluate the impact of time of year (summer heat, rate of degree day accumulation) separately from duration of infection period in defining the ‘time of inoculation’ parameter in vine recovery rate.

**Table 1.** Evidence supporting an effect of climate on winter recovery of X. *fastidiosa* infected grapevines.

| Potted or inground<sup>a</sup> | Field or chamber | Key Result | Reference |
| --- | --- | --- | --- |
| **Studies supporting a general effect of climate on spread of *Xf*** |  |  |  |
| Potted | Chamber | *Xf* populations growth in plants is temperature dependent | (Feil and Purcell 2001) |
| Potted | Chamber | Positive association of temperature with transmission efficiency | (Daugherty et al. 2009) |
| Potted | Chamber | Rates of pathogen transmission depended on climate | (Daugherty et al. 2017) |
| **Studies supporting an effect of inoculation date on winter recovery** |  |  |  |
| Inground | Field | Vines inoculated late in season more likely to recover than vines inoculated early in the season<br>Recovery differed with cultivar and location | (Purcell 1981) |
| Inground | Field | Vines inoculated late in summer more likely to recover than vines inoculated early in the season<br>Significant differences in recovery among locations | (Feil et al. 2003) |
| Potted | Field | Vines inoculated late in season more likely to recover | (Daugherty and Almeida 2019) |
| **Studies supporting an effect of cold period and duration on winter recovery** |  |  |  |
| Potted | Chamber | Results provided the basis for winter recovery hypothesis | (Purcell, A.H. 1977) |
| Potted | Field | Results suggest a non-linear relationship of winter temperature with recovery | (Purcell 1980) |
| Potted | Field | Rates of winter recovery varied with location and cultivar | (Lieth et al. 2011) |
| Potted | Chamber | Recovery rates increased with decreasing temperature, but varied with cultivar and year | (Lieth et al. 2011) |

<sup>a</sup> Some studies used potted vines held outdoors and some studies used established planted grapevines.

Regarding location, vineyard production in the state of California falls mainly within six distinct growing regions with quantifiably different climates (Fig. 1; North Coast, Sierra Foothills, Sacramento Valley, San Joaquin Valley, Central Coast, and Southern California). All of these areas are affected by Pierce’s disease although seasonality and severity of disease differ across regions, and vary by year (Vanhove et al. 2020; Sisterson et al. 2020; Haviland et al. 2021). Previous studies on winter recovery have largely focused on climates representative of grape growing regions in northern California (North Coast, Sierra Foothills, Sacramento Valley; Fig. 1). These regions produce high value wine grapes, but constitute a small portion of the overall grape production region. In contrast, little information is available regarding winter recovery of vines under climate conditions representative of southern and inland areas (Southern California, San Joaquin Valley), although approximately 85% of table grape and 75% of wine grape production takes place there. The limited observations of vine recovery (or lack of) that have been made in these areas suggest very different dynamics than more northern and coastal regions (Daugherty and Almeida 2019). In this study, temperature conditions were chosen to represent conditions typical of the San Joaquin Valley, both in regards to summer high temperature range, as well as winter chill hours.

**Figure 1.**
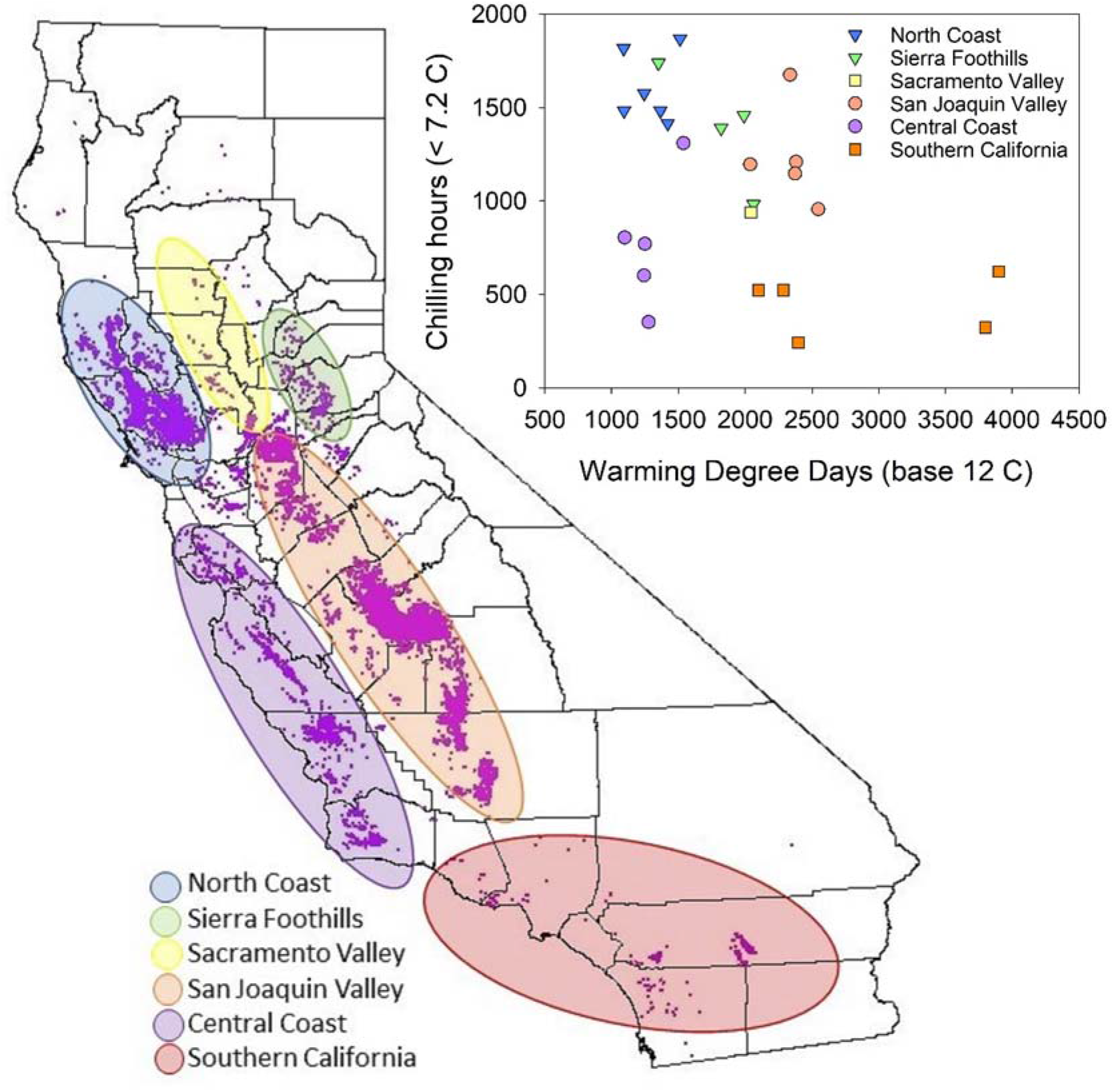
Location and climate associated with California’s five grape growing regions: North Coast, Sierra Foothills, Central Valley, Central Coast, and Southern California. To quantify summer and winter climate, weather station data was evaluated from sites within all regions (North Coast: Oakville, Knights Valley, Windsor, Santa Rosa, and Carneros; Sierra Foothills: Plymouth, Camino, Diamond Springs, and Auburn; Sacramento Valley: Davis; San Joaquin Valley: Parlier, Bakersfield, Porterville, Oakdale, and Orange Cove; Central Coast: Santa Ynez, Salinas, Lompoc, Nipomo, San Luis Obispo; Southern California: Temecula Valley (west), Temecula Valley (east), Coachella Valley, Escondido, and Thermal South). For summer climate degree days was determined with a base of 12 C. For winter climate, chill hours during dormancy were determined (hours < 7.2 C). Pixels in the map represent the distribution of grape plantings in California; for details on map creation see Sisterson et al. (2010).

In addition to time of inoculation and location, cultivar is the third prominent factor to consider in winter recovery dynamics. Within California, different types (wine, table and raisin) of grapes are grown with different genetic backgrounds and sensitivities to cold and *X. fastidiosa* (Rashed et al. 2011, 2013; Ferguson et al. 2014). More than 80 table grape and 60 wine grape cultivars are grown throughout the different production regions, many of which have not been evaluated for winter recovery rates. In general, table grapes are considered to be more susceptible to Pierce’s Disease (e.g. higher bacterial populations and more visible symptoms) than wine grapes, though studies comparing multiple cultivars are limited (Deyett et al. 2019; Rashed et al. 2011; Raju and Goheen 1981). Previous studies have reported that grape cultivars Scarlet Royal, French Colombard, and Thompson seedless have higher bacterial loads and symptoms compared to cultivars Cabernet Sauvignon, Crimson seedless and Merlot. It is likely that increased susceptibility would correspond to less recovery in most cases, as a systemic infection would develop more quickly in highly susceptible cultivars. Cabernet Sauvignon has been evaluated for winter recovery in several previous studies, but most cultivars for which experimental winter recovery data is available have only been included in a single study, often in only a single region (Table 1).

The overall goal of this study is to define a conceptual framework for understanding the impact of both summer and winter temperatures on grapevine recovery from *X. fastidiosa* infection. In addition, winter recovery of 6 grapevine cultivars under conditions representative of the San Joaquin Valley were evaluated to provide estimates of winter recovery for grape growing areas with hot summers and mild winters. To facilitate comparison with previous studies, cultivar Cabernet Sauvignon was included, along with two additional wine grape cultivars, Zinfandel and Sauvignon Blanc. ‘Zinfandel’ was previously found to be less susceptible to *X. fastidiosa* in greenhouse inoculation experiments than ‘Cabernet Sauvignon’, and ‘Sauvignon Blanc’ has intermediate susceptibility (Rashed et al. 2013; UC Statewide IPM Program (UC IPM)). Since table grape production is located in the San Joaquin Valley, three table grape cultivars were also included in this study: ‘Thompson seedless’, ‘Flame seedless’, and ‘Scarlet Royal’. Cultivar Thompson seedless is considered to be highly susceptible *to X. fastidiosa* infection, similar to ‘Flame seedless’ and ‘Scarlet Royal’ (Rashed et al. 2013; Deyett et al. 2019). The results of this study will facilitate improved risk models across a wider range of climate conditions, to better predict Pierce’s disease risk in new geographic areas and under future climate scenarios.

## Methods

### Conceptual model

To evaluate the impact of a range of climate conditions on disease, and to provide context for experimental results, a conceptual model was developed based on the known factors influencing disease severity and winter recovery (inoculation date, climate, and cultivar susceptibility). On inoculation, bacteria are only present at the inoculation site. Overtime, the *X. fastidiosa* population increases and spreads through the plant (Hill 1995). Because *X. fastidiosa* population growth (Feil and Purcell 2001) and onset of Pierce’s disease symptoms (Daugherty et al. 2017) are positively associated with temperature, rate of infection progression of *X. fastidiosa* in grapevines is greater in warm compared to cool climates. Conceptually, the *X. fastidiosa* infection status of a vine (represented here as N_xf_) is a parameter that includes bacterial population in the plant as well as symptom progression. Symptom progression being influenced by plant immune responses in addition to bacterial replication within the xylem (Rapicavoli et al. 2018; Abou Kubaa et al. 2019). On inoculation, *N_Xf_* is near zero and increases over time as a function of time and temperature (Fig. 2A). Infection status in susceptible vines increases until a maximum level of infection 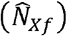 is reached or vines enter dormancy (Fig. 2A). Bacteria are expected to replicate faster at higher temperatures, and although not well characterized in grapevine-*X. fastidiosa* interactions, plant immune responses are known to be influenced by higher temperatures as well (Cheng et al. 2013). Importantly, *X. fastidiosa* populations are known to increase and decrease during the season in chronically infected vines in the field (Sisterson et al. 2020). Both size of *X. fastidiosa* populations in vines, and also symptom severity are known to vary among cultivars (Rashed et al. 2011, 2013). Accordingly, *X. fastidiosa* equilibrium infection status 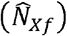 is cultivar dependent and the magnitude of 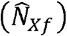 may increase and decrease throughout the year.

**Figure 2.**
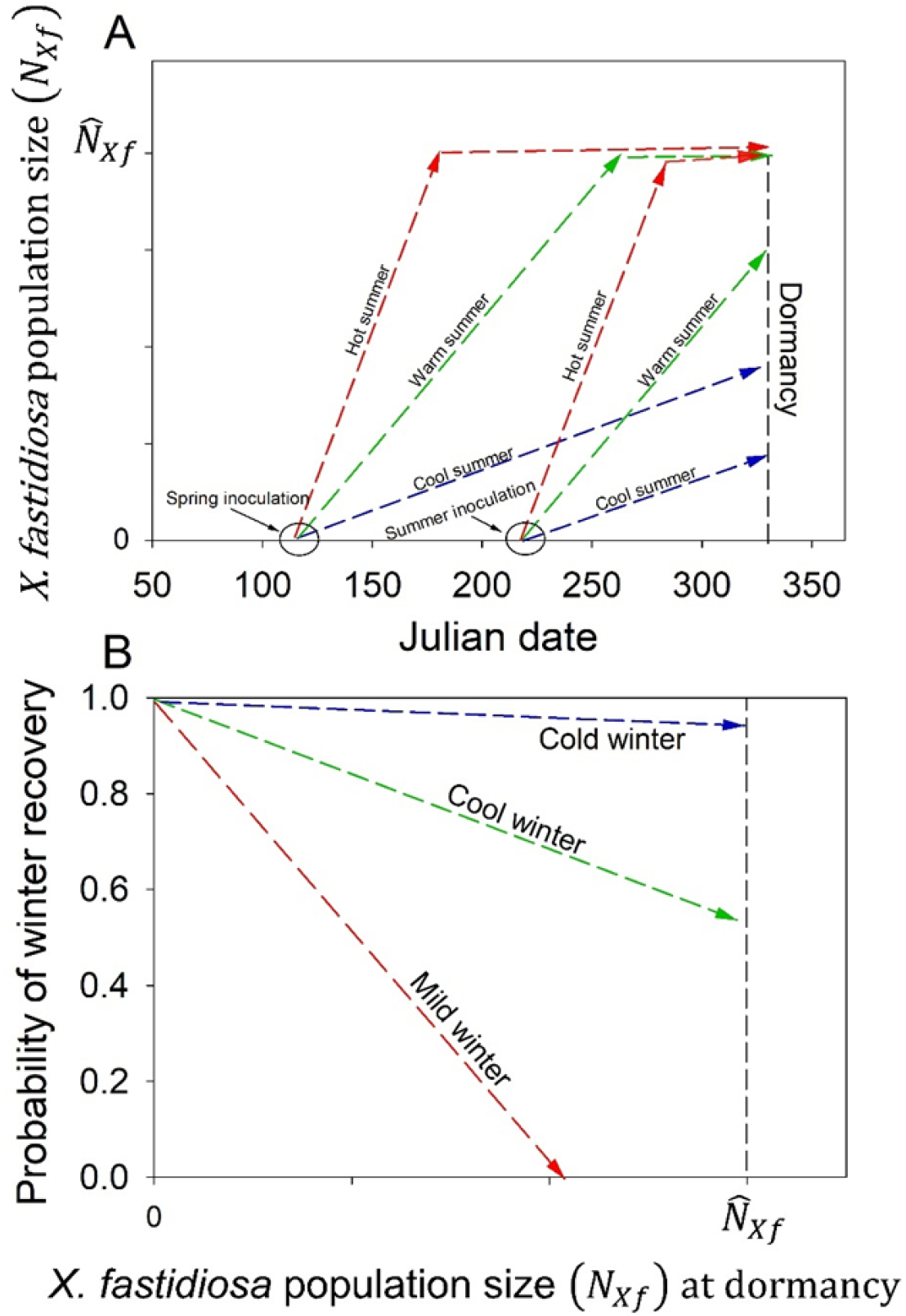
Conceptual model of winter recovery. A) The rate *of X. fastidiosa* population growth in a vine is temperature dependent. Accordingly, size of the *X. fastidosa* population in a vine at dormancy depends on the date the vine was inoculated and on summer climate, with populations increasing faster in warm compared to cool climates. B) Rates of winter recovery depend on winter climate and size of the *X. fastidiosa* population at the time vines enter dormancy. Probability of winter recovery is presumed to decline as winter temperatures decrease and as size of the *X. fastidiosa* population at dormancy increases. While evidence (Table 1) supports the directionally of the relationships described in panels A and B, the shape of the functions are unknown and may be non-linear.

As vines enter dormancy, the *X. fastidiosa* infection status of vines inoculated during early summer may have reached 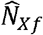, whereas *X. fastidiosa* infection status of vines inoculated in late summer may not have reached 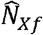 prior to dormancy. Probability of winter recovery depends on *X. fastidiosa* infection status (*N_Xf_*) at dormancy and winter climate (Fig. 2B). It is not currently clear if this is influenced more by symptom severity or bacterial population or equally by both. Vines with low values for *N_Xf_* are presumed to require less exposure to cold temperatures to recover than vines with high values for *N_Xf_*. While longer durations at cooler temperatures favor winter recovery (Table 1), the precise conditions required for winter recovery are poorly defined, and require additional experimental testing.

### Relative likelihood of winter recovery in each of California’s grape growing regions

Based on the conceptual model (Fig. 2A, B), winter recovery of vines is least likely in climates with hot summers and mild winters and most likely in climates with cool summers and cool winters. Thus, categorization of the summer and winter climates of each of the grape growing regions in California provides a method for establishing relative likelihood of winter recovery.

To quantify the summer and winter climate of each of California’s grape growing regions, weather station data was evaluated from sites within all regions for the period from 2018 thru 2021 (North Coast: Oakville, Knights Valley, Windsor, Santa Rosa, and Carneros; Sierra Foothills: Plymouth, Camino, Diamond Springs, and Auburn; San Joaquin Valley: Parlier, Bakersfield, Porterville, Oakdale, Orange Cover; Sacramento Valley: Davis; Central Coast: Santa Ynez, Salinas, Lompoc, Nipomo, San Luis Obispo; Southern California: Temecula Valley (west), Temecula Valley (east), Coachella Valley, Escondido, and Thermal South). For winter climate, chill hours (hours < 7.2 °C) were determined for each location and year using the University of California Fruit and Nut crop information center (https://fruitsandnuts.ucanr.edu/Weather_Services/Chill_Calculators/). For summer climate, degree days with a base of 12 °C were calculated (McMaster and Wilhelm 1997) for each location and each year. Degree days were calculated using summer temperature data for each location acquired from the University of California Statewide Integrated Pest Management Program website (http://ipm.ucanr.edu/WEATHER/index.html). A base of 12 °C was used as temperatures must be > 12 °C for *X. fastidiosa* populations to grow (Feil and Purcell 2001). For each measure, the average was determined over the period from 2018 thru 2021.

### Plants, bacteria, and inoculations

One-year-old potted grapevines (*Vitis vinifera*) were grown in a greenhouse with artificial light to maintain 16 hr light/8 hr dark year-round and semi-temperature controlled to maintain temperature over 18.3°C. Inoculation experiments were done twice (2018 and 2020). Vines were inoculated using a previously described needle inoculation protocol (Roper et al. 2007) with *X. fastidiosa* Stag’s Leap on the first day of the warming period (25-Jun-18, 15-Jun-20, and 19-Aug-20). Inoculation dose consisted of 40 μl total bacterial cell suspension with a concentration of 10^8^ cfu/ml per vine. For each cultivar by warming period combination, 15 vines were inoculated. Five mock-inoculated vines were used as negative controls. Vines were held in the greenhouse with position randomized with regards to cultivar and treatment combination. After inoculation, vines were exposed to one of three warming periods in the greenhouse designed to represent different seasonal inoculation dates (Fig. 4). Two of the warming periods lasted 8 weeks, but one was from late June thru mid-August (2018: 25-Jun to 21-Aug; 2020: 15-Jun to 13-Aug) and the other was from mid-August through mid-October (2020 only: 19-Aug to 9-Oct). A final 16-week warming period treatment covering the full range of both 8 week warming periods was also evaluated (2018: 25-Jun to 16-Oct; 2020:15-Jun to 9-Oct). Temperature readings were collected from the greenhouse every 15 minutes for the entire duration of the experiments using a HOBO data logger device. Subsequently, heat unit accumulation (degree days with a base of 12 °C) was determined for each warming period. Three cultivars were evaluated in 2018: Cabernet Sauvignon, Scarlet Royal, and Zinfandel. In 2018, only two of the three warming periods were evaluated (25-Jun-18 to 21-Aug-18 and 25-Jun-18 to 16-Oct-18). In 2020, all three warming periods were evaluated, and number of cultivars evaluated increased to six (Cabernet Sauvignon, Sauvignon Blanc, Thompson Seedless, Flame Seedless, Scarlett Royal, and Zinfandel). Cultivars Scarlet Royal and Flame Seedless were own-rooted and the rest of the cultivars were grafted on the following rootstocks: Cabernet Sauvignon on 110R, Sauvignon Blanc on Schwarzmann, Thompson Seedless on Salt Creek, Zinfandel on SO4. All plants were grown in 2-gallon pots in Sunshine mix #1 (SunGro Horticulture). After the warming period, vines were pruned to a single cane and placed directly in cold chambers for 5 weeks at 4.4 °C. This cold treatment resulted in an accumulation of 840 chill hours; typical of the long-term trend for the southern San Joaquin Valley area (Arvin, CA Fig. 3) where two epidemics of Pierce’s disease have occurred approximately a decade apart (Tubajika et al. 2004; Sisterson et al. 2020). Chill hours accumulated in the San Joaquin Valley area are less than those observed on the North Coast, but greater than those observed in Southern California and on the Central Coast (Fig. 1).

**Figure 3.**
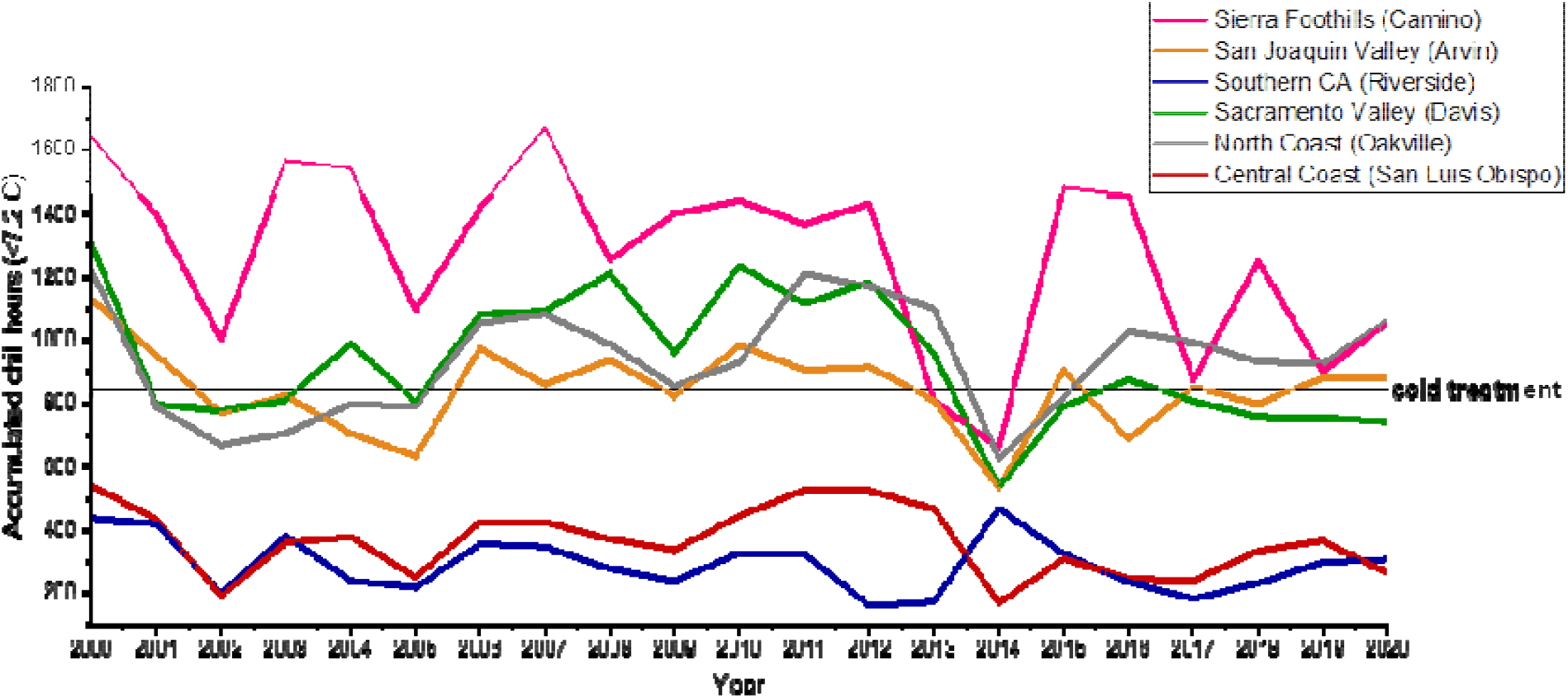
Historical chill hours observed in different grape growing regions of California. Accumulated chill hours (hours < 7.2 °C) were determined for each location and year using the University of California Fruit and Nut crop information center (https://fruitsandnuts.ucanr.edu/Weather_Services/Chill_Calculators/). For cold treatment of infected vines, a chill hour period similar to the San Joaquin Valley (Arvin, CA) was selected to represent grape growing regions with hot summers that have experienced severe Pierce’s disease epidemics.

**Figure 4.**
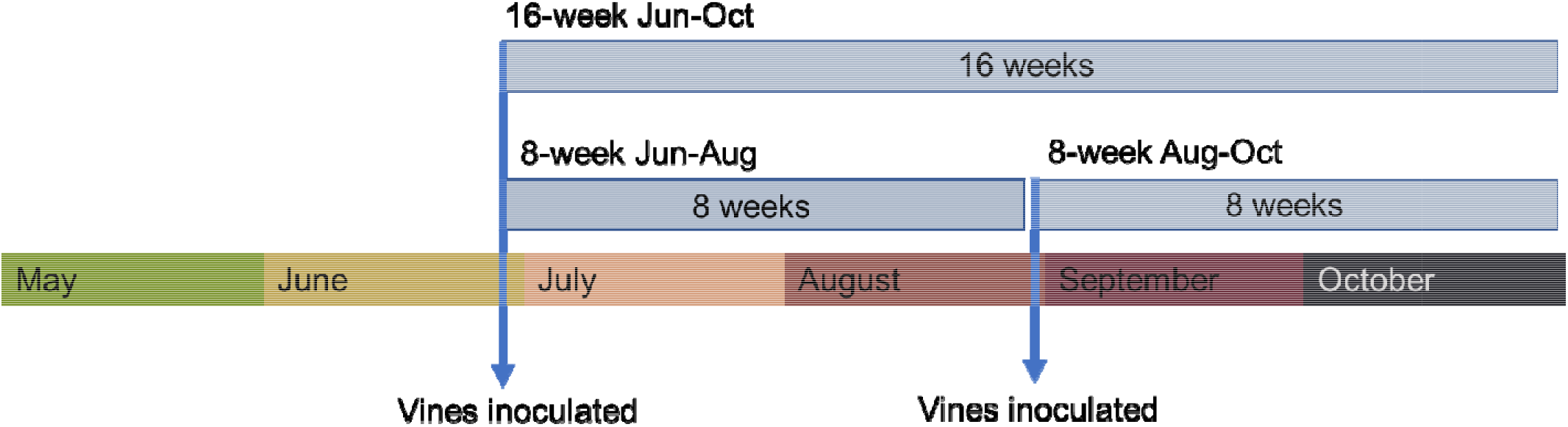
Experimental set-up for inoculation and cold treatment experiments. Plants were separated into three different treatment groups. Two groups were inoculated in late June, one of which was transferred to the cold chamber after 8 weeks (8-week Jun to Aug) and the other after 16 weeks (16-week Jun to Oct). The third group was inoculated in late August and transferred to the cold chamber after 8 weeks (8-week Aug to Oct). All the plants were kept in the same greenhouse between inoculation and cold treatment where temperature was semi-controlled but subject to daily fluctuations depending on weather conditions. Cold treatment consisted of 5 weeks at 4.4C for all treatments.

Leaf petioles were collected from all vines and subjected to qPCR for *X. fastidiosa* detection immediately prior to cold treatment and again 20 weeks after the end of the cold treatment when plants had been able to re-grow fully and infections had time to become re-established. Similarly, disease symptoms were evaluated immediately prior to cold treatment and again 20 weeks after the end of the cold period. Symptom evaluations followed a previously described 0-5 rating scale and were conducted blind to the treatment (Roper et al. 2007).

### Quantitative PCR testing

For qPCR testing, three petioles were collected from each vine and pooled for DNA extraction. Petiole samples were lyophilized and ground to fine powder using a TissueLyzer II bead mill (Qiagen). DNA was extracted from ground tissue using a CTAB/phenol/chloroform extraction protocol as previously described (Wei et al. 2021). DNA samples were quantified with a Quant-IT DNA assay kit (ThermoFisher Scientific) and diluted 1:10 in sterile water for PCR. Quantitative PCR was performed as previously described (Wei et al. 2021) with AB Fast SYBR Green Master Mix (ThermoFisher Scientific) and primers XfITS145-60F 5’-TACATCGGAATCTACCTTATCGTG-3’ and XfITS145-60R 5’-ATGCGGTATTTAGCGTAAGTTTC-3’ (Ledbetter and Rogers 2009).

### Analyses

A generalized linear model with a logit link function and binomial error was used to test effects of cultivar, warming treatment, and the interaction between cultivar and warming treatment on probability that a vine tested positive at the end of the warming treatment. Contrasts were used to compare cultivars within warming treatments. An Analysis of Variance was used to test effects of cultivar, warming treatment, and the interaction of warming treatment on symptom expression at the end of the warming period. If overwinter recovery occurs, fewer plants should test positive for *X. fatsidiosa* after the cold treatment than before. To determine if recovery occurred, the difference in the proportion of plants infected before and after the cold treatment was determined:

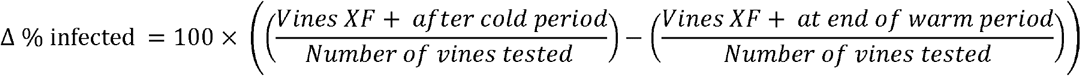

If the change is positive, number of infected plants increased after the cold treatment suggesting no winter recovery. In contrast, if the change is negative, number of infected plants decreased after the cold treatment suggesting winter recovery occurred. To determine if rates of winter recovery were significant, 95% confidence intervals were constructed for each cultivar by warming period combination assuming binomial error. If the mean was negative and the confidence interval did not include zero, recovery was considered significant. Vines that died during the cold treatment were omitted from analysis (8 out of 360 vines).

## Results

### Relative likelihood of winter recovery in California’s grape growing regions is dependent on summer and winter conditions

Across the state of California there is considerable variation in climate among grape growing regions (Fig. 1). Accumulation of degree days varied 3.6-fold between locations with the warmest and coolest summers. Likewise, accumulation of chill hours varied 7.7-fold between the location with the coldest and warmest winter. In regions such as the North Coast with relatively cool summers and cool winters (Fig 1), progression of *X. fastidiosa* infection in vines is likely to be slow with *X. fastidiosa*-infected vines often not reaching 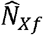 prior to dormancy (e.g., Fig. 2A). Thus, on the North Coast, winter recovery rates are likely to be high as values of *N_Xf_* at dormancy are likely to be low and winters are cool. In fact, winter recovery studies have focused on climates of the North Coast and Sierra foothills (Table 2); regions of California likely to experience the highest rates of vine recovery.

In regions with hot summers such as Southern California and the San Joaquin Valley (Fig. 1), infection progression of *X. fastidiosa* populations in vines is likely to be rapid, with vines often achieving values of 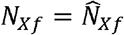 prior to dormancy (e.g. Fig. 2A). While degree day accumulation during the summer is high in Southern California and in the San Joaquin Valley, winter climates differ between the regions. Winters in the San Joaquin Valley are cooler than winters in Southern California. Chill hour accumulation in portions of the San Joaquin and Sacramento Valleys are as high as on the North Coast (Fig. 1). Thus, some degree of winter recovery is likely to be observed in portions of these inland areas that experience cooler winters (Purcell 1981; Feil et al. 2003; Daugherty and Almeida 2019). Given the hot summers and mild winters observed in Southern California and the southern portion of the San Joaquin Valley, winter recovery may be expected to be limited in these areas. Currently, information on winter recovery is available for only 9 wine grape cultivars and 2 table grape cultivars, with only 2 cultivars evaluated in more than one region (Table 2). It is likely that susceptibility of a specific cultivar will impact the amount of degree days required to achieve 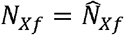 and therefore the rate of recovery under different climate scenarios.

**Table 2.** Cultivars and regional location of studies conducted on winter recovery.

| Cultivar | North Coast<sup>a</sup> | Sierra Foothills<sup>a</sup> | Sacramento Valley<sup>a</sup> | San Joaquin Valley | Central Coast<sup>a</sup> | Southern California<sup>a</sup> | % of total area planted in CA<sup>b</sup> |
| --- | --- | --- | --- | --- | --- | --- | --- |
| **Red wine type grapes** |  |  |  |  |  |  |  |
| Barbera | x | x | x | Purcell 1981<sup>c</sup> | x | x | 0.6 |
| Cabernet Sauvignon | Purcell 1981<br>Feil et al. 2003<br>Lieth et al. 2011 | Lieth et al. 2011 | Feil. et al. 2003<br>Lieth et al. 2011 | x | x | x | 10.6 |
| Ruby Cabernet | x | x | x | Purcell 1981<sup>c</sup><br>Feil et al. 2003<sup>c</sup> | x | x | 1.2 |
| Pinot noir | Lieth et al. 2011 | Lieth et al. 2011 | Lieth et al. 2011<sup>c</sup> | x | x | x | 5.4 |
| Merlot | Feil et al. 2003 | x | x | x | x | x | 4.1 |
| **White wine type grapes** |  |  |  |  |  |  |  |
| Chardonnay | Purcell 1981<br>Feil et al. 2003 | x | x | x | x | x | 10.3 |
| Chenin blanc | Purcell 1981 | x | x | x | x | x | 0.5 |
| Flora | Purcell 1981 | x | x | x | x | x | <0.1 |
| White Reisling | Purcell 1981 | x | x | x | x | x | 0.4 |
| **Raisin and table type grapes** |  |  |  |  |  |  |  |
| Red globe | x | x | x | Daugherty and Almeida 2019<sup>d</sup> | x | x | 0.7 |
| Thompson seedless | x | x | x | Daugherty and Almeida 2019<sup>d</sup> | x | x | 13.6 |
|  |  |  |  |  |  | Total % | 47.5 |

<sup>a</sup>. An “x” indicates that results are not available for that region
<sup>b</sup>. Estimates of area planted from CDFA (2020).
<sup>c</sup>. Field sites were located >100 miles north of areas impacted by the glassy-winged sharpshooter.
<sup>d</sup>. Field sites were located in a glassy-winged sharpshooter infested area.

### Degree-days accumulated during experimental warming treatments represent different inoculation dates in the San Joaquin Valley

In greenhouse experiments, the number of degree days accumulated in the two 8-week warming treatments was similar, although more degree days were accumulated during the 8-week period between June and August than in the 8-week period between August and October (Fig. 5). Number of degree days accumulated during the 16-week warming treatment was double the number accumulated during the 8-week treatments (Fig. 5).

**Figure 5.**
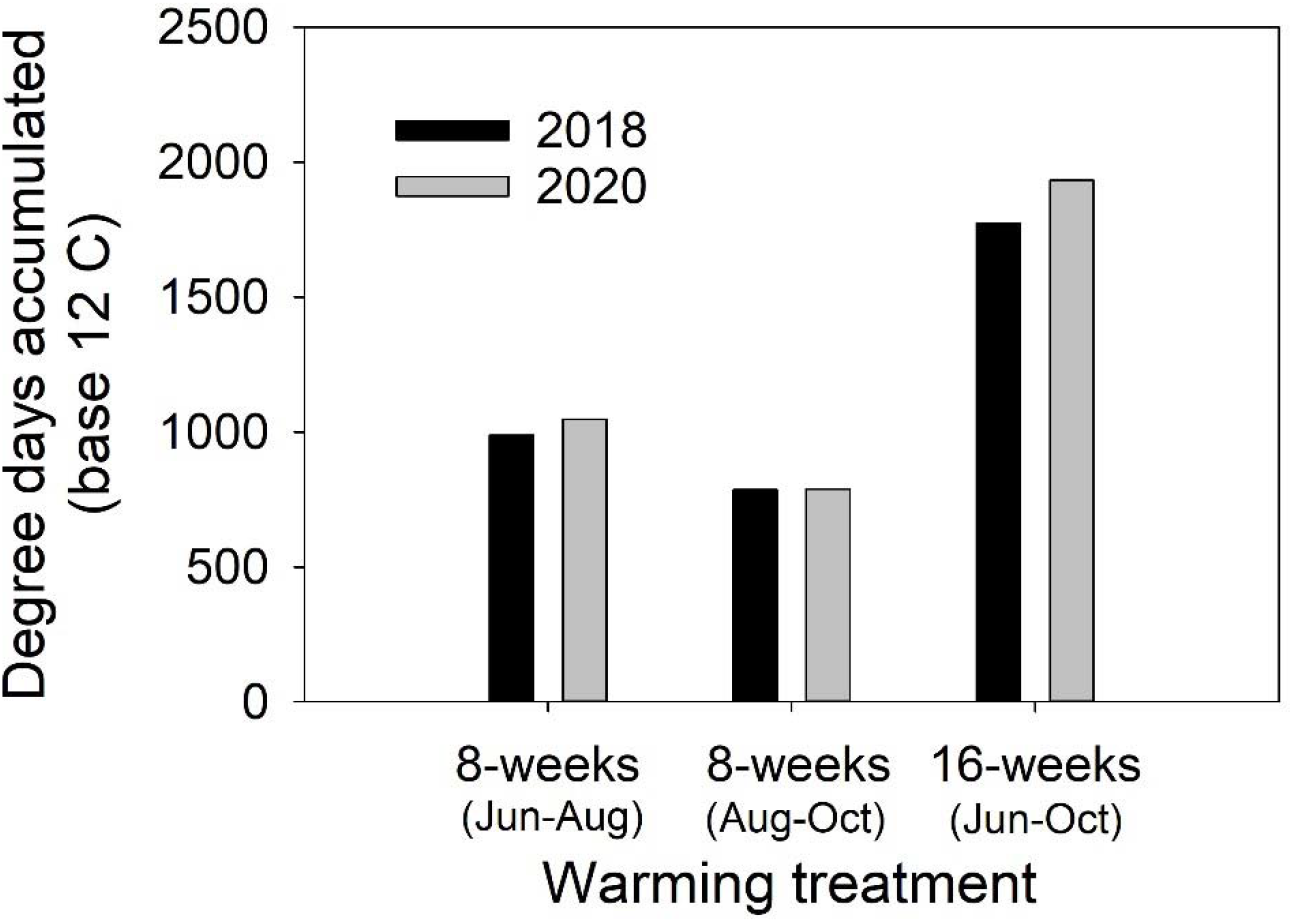
Number of degree days accumulated for each warming treatment by year combination. Temperature data was collected in the greenhouse every 15 minutes for the duration of the warming period using a HOBO Data Logger. Degree days were calculated using a base of 12 C. Experiment was repeated in two separate years, 2018 and 2020.

By comparing number of degree days accumulated in the greenhouse prior to cold treatment to historical temperature records from the southern San Joaquin Valley (Arvin, CA), estimates of equivalent inoculation dates in that location may be determined. As temperatures in the greenhouse were similar to summer temperatures in the San Joaquin Valley, the number of degree days accumulated during the 8-week warming period from mid-August through mid-October represented an inoculation date in Arvin, CA of mid-August. The number of degree days accumulated during the 8-week warming period from mid-June through mid-August represented an inoculation date in Arvin of early August. Finally, the 16-week warming period represented an inoculation date in Arvin of mid-June.

### Effects of cultivar and warming treatment on infection status prior to cold treatment

For vines inoculated in 2018, there was a significant effect of cultivar (χ^2^ =10.6, df = 2, P = 0.005) and the interaction of cultivar with warming treatment (χ^2^ = 7.6, df = 2, P = 0.02), but no main effect of warming treatment (χ^2^ = 2.8, df = 1, P = 0.09) on proportion of vines testing positive via qPCR at the end of the warming treatment (Fig. 6A). For vines inoculated in 2020, there was a significant effect of cultivar (χ^2^= 11.2, df = 5, P = 0.048), but no effect of warming treatment (χ^2^= 4.3, df = 2, P = 0.11) or the interaction between cultivar and warming treatment (χ^2^= 16.5, df = 10, P = 0.085) on proportion of vines infected at the end of the warming treatment (Fig. 7A). As there was no effect of warming treatment on proportion of vines testing positive, the degree days required for *X. fastidiosa* populations to achieve a value of 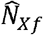 was less than that accumulated for the lowest degree day treatment tested (785-degree days).

**Figure 6.**
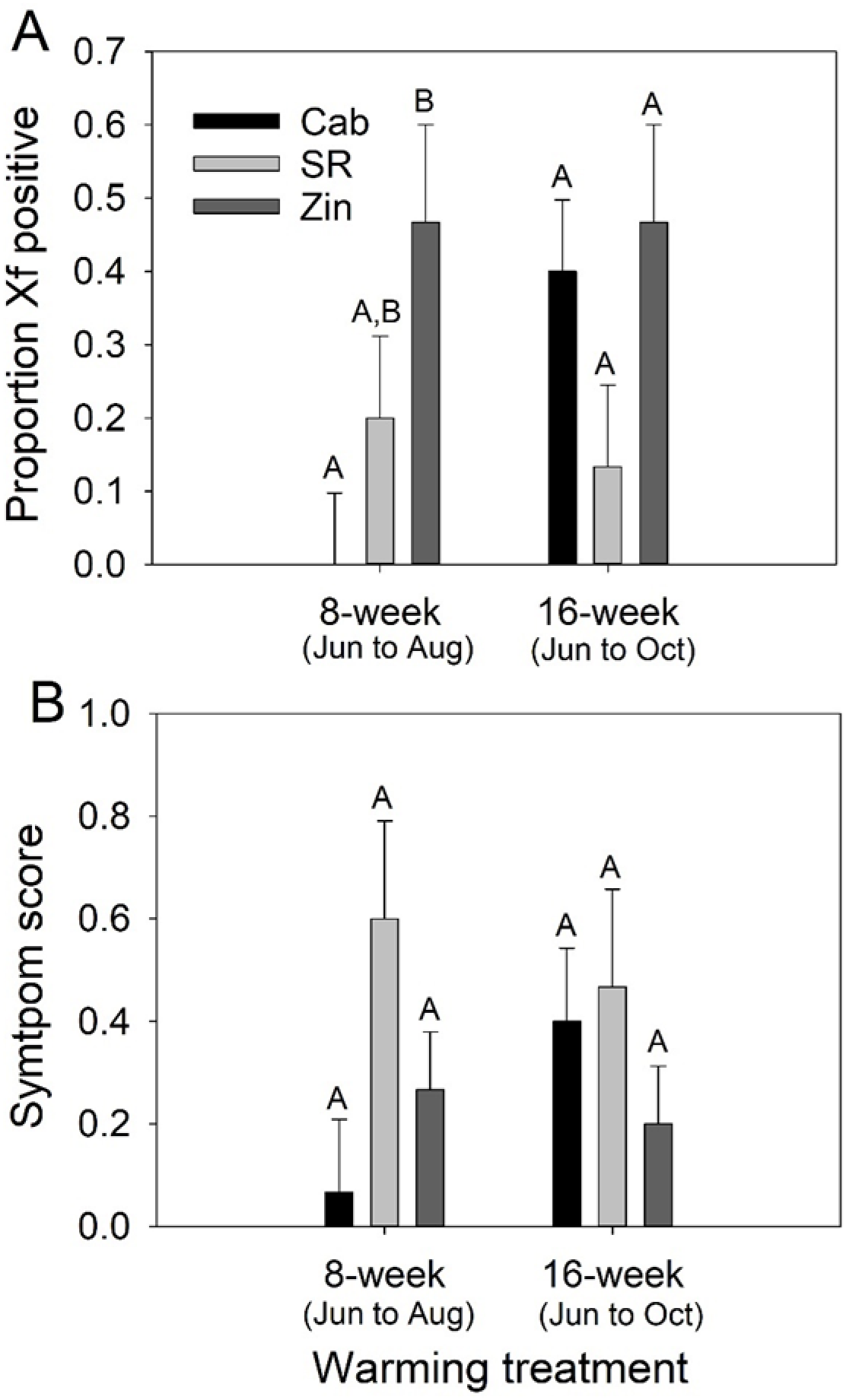
Infection status of vines at the end of warming treatments in 2018. A) Proportion (± SE) of plants positive for *X. fastidiosa* via qPCR for each cultivar and warming treatment combination. B) Mean (± SE) symptom score for each cultivar and warming treatment combination. Different letters above bars indicates significant difference among cultivars within warming treatment. Cab = Cabernet Sauvignon, SR = Scarlet Royal, Zin – Zinfandel

**Figure 7.**
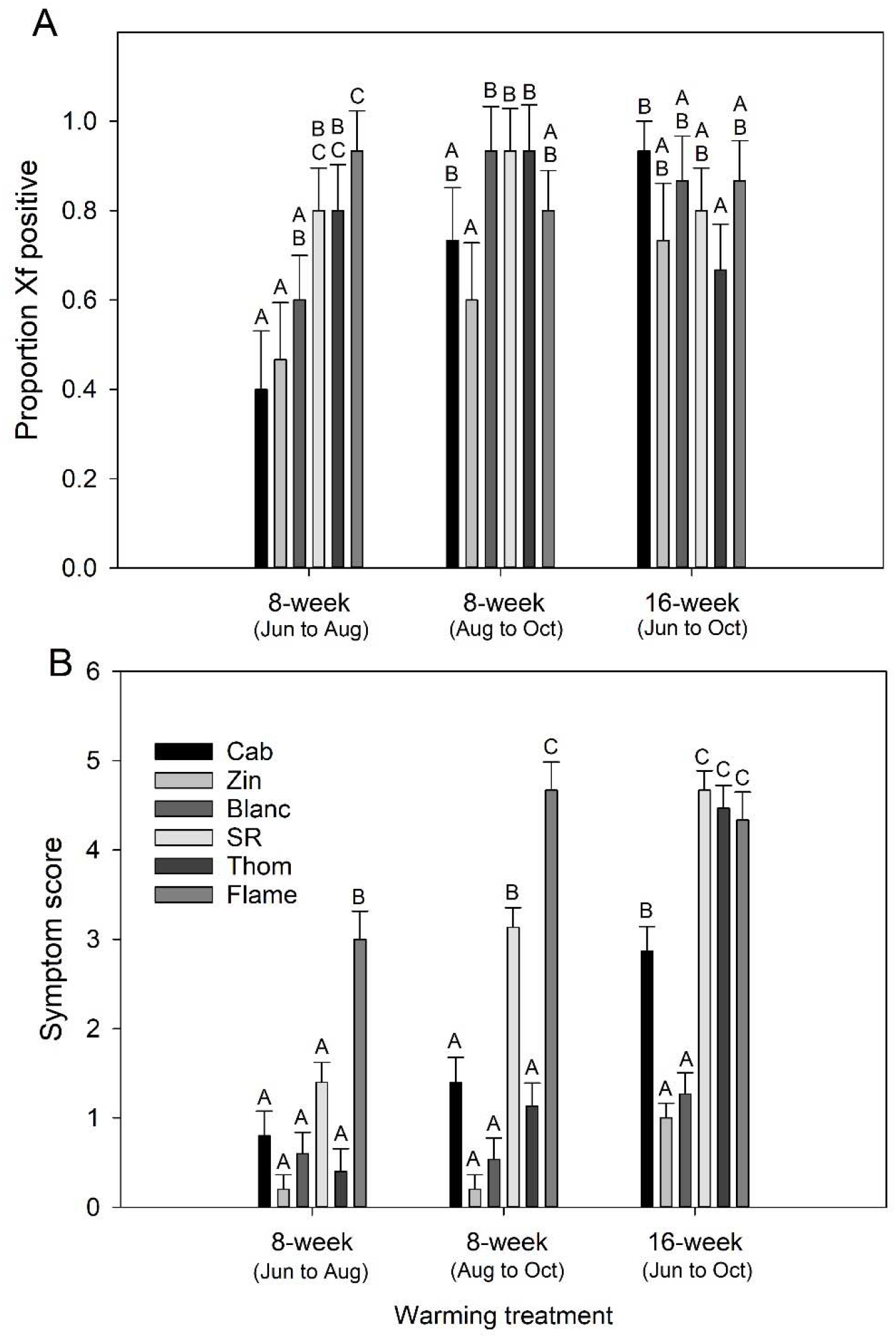
Status of vines at the end of warming treatments in 2020. A) Proportion (± SE) of plants positive for *X. fastidiosa* via qPCR for each cultivar and warming treatment combination. B) Mean (± SE) symptom score for each cultivar and warming treatment combination. Different letters above bars indicates significant difference among cultivars within warming treatment. Status of vines at the end of warming treatments in 2018. A) Proportion (± SE) of plants positive for *X. fastidiosa* via qPCR for each cultivar and warming treatment combination. B) Mean (± SE) symptom score for each cultivar and warming treatment combination. Different letters above bars indicates significant difference among cultivars within warming treatment. Cab = Cabernet Sauvignon, Zin = Zinfandel, Blanc = Sauvignon Blanc, SR = Scarlet Royal, Thom = Thompson seedless, Flame = Flame seedless

For vines inoculated in 2018, there was no effect of cultivar, warming treatment, or the interaction of warming treatment with cultivar on symptom severity (Fig. 6B, whole model: F = 1.6, df = 5, 84, P = 0.17). There was a significant effect of cultivar (F = 87.0, df = 5, 252, P < 0.0001), warming treatment (F = 101.5, df = 2, 252, P < 0.0001), and the interaction of warming treatment with cultivar on symptom severity (F = 10.73, df = 10, 252, P < 0.0001). In 2020, vines held under warm conditions for 16 weeks had significantly greater symptom expression then vines held under warm conditions for 8 weeks (Fig. 7B, F = 173.8, DF = 1, 252, P < 0.0001).

### Effect of warming treatment and cultivar on winter recovery

Overall, winter recovery was limited and not clearly related to a single warming treatment (Fig. 8). In 2018, no significant recovery was observed for the 3 cultivar and two warming treatments evaluated. In 2020, significant recovery was observed for 5 of 18 cultivar by warming period combinations. The cultivar Zinfandel displayed significant recovery across all three warming period treatments (Fig. 8B). Results suggest winter recovery sometimes occurs in climates with hot summers and milder winters representative of inland and southern California grape growing regions, but is limited and likely cultivar dependent.

**Figure 8.**
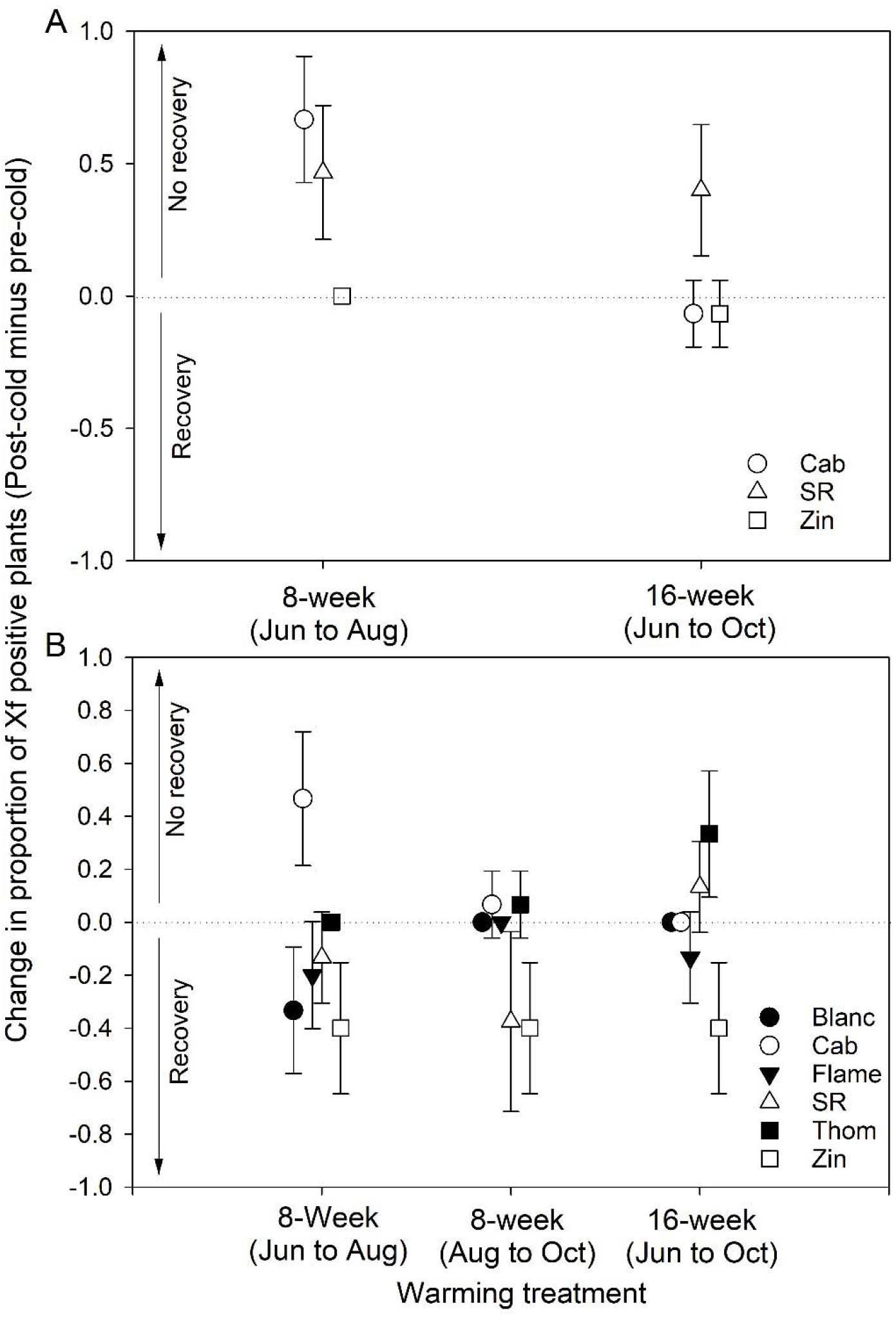
Change in proportion of positive plants (post cold minus pre cold) in 2018 (A) and 2020 (B). Bars represent 95% confidence intervals. Curing is considered significant if the change in the proportion of infected plants is negative and the bars do not overlap with zero.

## Discussion

Epidemiology of insect transmitted plant pathogens varies among crop growing regions for a variety of factors including climate. Understanding the specific impacts of temperature on plant disease is essential for transferring knowledge across different regions, and for predicting disease dynamics under future climate scenarios (Chaloner et al. 2021). Pierce’s disease effects grape production in tropical and subtropical regions throughout the world, but with highly variable epidemic severity depending on regional differences in climate, insect vectors, and vineyard management practices (EPPO Global Database, 2022). Based on the current known climate suitability of *X. fastidiosa* there is high potential for new outbreaks to occur if the pathogen and/or its vectors are introduced into new regions (Godefroid et al. 2019). In addition, there is risk of increased disease severity and crop impacts in areas where *X. fastidiosa* is endemic due to new vector introductions, changes in management practices, or global warming (Haviland et al. 2021). Since grapevines are also cultivated in temperate areas with colder winters where *X. fastidiosa* is not currently found, expansion of the suitable range for Pierce’s disease is likely as temperatures increase in the future (Wallingford et al. 2007; Daugherty et al. 2017; Godefroid et al. 2019; Castex et al. 2018; Koch and Oehl 2018; Giménez-Romero et al. 2022; Godefroid et al. 2022). To better predict the impact of climate change on disease epidemiology and pathogen geographic range, it is necessary to conceptualize and empirically test specific parameters influencing disease severity and pathogen spread. Pierce’s disease epidemiology in California presents an ideal case study for characterizing climate factors associated with disease due to cultivation of grapevines across diverse climatic regions within the state, and a long history of research in the area on *X. fastidiosa* and its insect vectors.

Field trials are often conducted across a limited portion of a pathogens range, and for winter recovery of grapevines from Pierce’s disease, previous studies centered around northern and coastal areas of California (Table 2). In this study we focused on the impact of temperatures representative of inland and southern regions with hot summers and milder winters, to provide a more comprehensive picture of climate influence on winter recovery and to facilitate comparison with other regions worldwide and under global warming scenarios. To evaluate relative likelihood of winter recovery in California’s grape growing regions, summer and winter climates in each area were quantified and predictions developed based on a conceptual model. Given the cool summers and cool winters experienced on the North Coast (Fig. 1), winter recovery is likely to play an important role in Pierce’s disease epidemiology in that area. In contrast, with the hot summers and mild winters associated in southern California and the San Joaquin Valley, winter recovery is likely to be limited. Experimentally observed recovery rates of inoculated grapevines grown under specific temperature regimes supported minimal recovery with climate conditions representative of inland and southern grape growing regions in California (Arvin, CA)(Fig. 8). As indicated by the model of Daugherty and Almeida (2019), epidemics progress more rapidly in the absence of winter recovery. Since minimal recovery would be expected in inland and southern grape-growing regions in California, as well as other parts of the world with comparable climate conditions, winter recovery should not be viewed as a major factor affecting Pierce’s disease epidemiology in those areas. Predicted vine recovery rates also impact vineyard management practices such as insecticide treatments to reduce vector populations. In areas where recovery rates are higher, controlling vector populations late in the season may be unnecessary as late season infections will recover over the winter (Feil et al. 2003). In contrast, for areas with hot summers where recovery is expected to be minimal, vector control late in the season may be more important to prevent additional pathogen spread.

Several studies demonstrate an effect of inoculation date on likelihood of winter recovery (Table 1). To facilitate making broader predictions, future studies should report number of heat units accumulated prior to dormancy along with inoculation date. Vines held for 8-weeks under summer conditions for an accumulation of 785-degree days had similar rates of infection prior to cold treatments (Fig. 6A, 7A) and rates of winter recovery (Fig. 8) as vines held for 16-weeks under summer conditions for an accumulation of 1774-degree days. Accordingly, number of degree days required to achieve 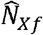 for most of the cultivars evaluated here is less than 785-degree days. Given typical summer temperatures in warmer growing regions, *X. fastidiosa* populations will likely reach 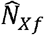 prior to dormancy provided vines are inoculated before September. In agreement, Daugherty and Almeida (2019) observed no winter recovery for vines inoculated in Arvin, CA with ~850-degree days remaining in the year (16-Jul in Arvin, CA) and limited recovery (~40%) for vines inoculated with ~550-degree days remaining in the year (3-Sep in Arvin, CA).

Implicit in the observation of an effect of inoculation date on winter recovery (Table 1) is that there are quantifiably different attributes associated with the *X. fastidiosa* infection progression in vines based on inoculation date. Here, the nebulous attribute of *X. fastidiosa* infection status of a vine was conceptually represented by the parameter *N_Xf_* which was presumed to increase overtime eventually reaching an equilibrium 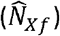 provided there were sufficient heat units remaining in the year prior to dormancy. While it is reasonable to presume that a vine inoculated in spring had more time for *X. fastidiosa* populations to spread and symptoms to develop than a vine inoculated in late summer, it is not readily clear how to quantify this attribute in field or laboratory tests. Distribution of *X. fastidiosa* in vines is variable and measurement from one or only a few leaves is likely to be insufficient to properly characterize the *X. fastidiosa* population in a whole plant. In addition, symptoms may not always correlate directly with bacterial population depending on the grape cultivar and it is unclear whether high bacterial population, advanced symptoms, or both drive the likelihood of winter recovery (Krivanek and Walker 2005; Krivanek et al. 2005; Riaz et al. 2020; Gambetta et al. 2007). It is possible that movement of *X. fastidiosa* to specific plant parts, or internal damage and obstruction of the xylem vessels is also critical to whether vines can recover. Plants infected with *X. fastidiosa* produce tyloses within the xylem vessels in response to presence of the pathogen causing vascular obstruction or xylem cavitation and contributing to the disease symptoms (Sun et al. 2013; Sabella et al. 2019; Ingel et al. 2021). The magnitude of this pathogen-induced response as well as variation in the xylem structure between cultivars can influence symptom severity (Sabella et al. 2019; Deyett et al. 2019). Since tylose formation is irreversible, severe xylem vessel obstruction could impact vine health into the next season and reduce likelihood of vine recovery. Other factors such as age of the plants may also play a role, as younger plants may be more susceptible to infection (Purcell 1981).

Previous studies (Tables 1 and 2) and this study (Fig. 6 and 7) report an effect of cultivar on winter recovery. To date information on winter recovery is available for only 9 wine grape cultivars and 2 table grape cultivars (Table 2). However, ~75 wine grape cultivars and ~80 table/raisin grape cultivars are grown in California (CDFA 2020), and globally the varietal diversity is much greater. Accordingly, no information regarding winter recovery is available for most cultivars grown worldwide. Lack of cultivar specific knowledge is compounded by an absence of information regarding effect of rootstock on winter recovery, or other compounding factors such as drought stress. Future field studies should attempt to include regionally popular cultivars in assessments, and further research is needed to define specific physiological factors that could be used to predict recovery as well as overall disease susceptibility. As ability to separate grapevines into genetic groups based on genomic comparisons improves (Bacilieri et al. 2013; Emanuelli et al. 2013; Riaz et al. 2018; Riaz et al., 2020) it may be possible to identify genetic groupings associated with a greater degree of winter recovery than other groupings.

Winter temperature regimes required to observe winter recovery are poorly defined. In the cold chamber studies conducted here limited curing was observed for vines held for 5-weeks at 4.4 C, resulting in an accumulation of 840 chill hours which is typical for the San Joaquin Valley area (Fig. 3). The chambers were set to maintain constant temperature. It is unknown to what extent fluctuating temperatures affect winter recovery. It is possible that brief periods of unusually low temperature could play an important role in winter recovery. For example, initial observation on winter recovery (Purcell 1974) used cold chambers set to very low temperatures (−8 C and −12 C) for periods measured in hours. The current conceptual model on winter recovery presumes that colder temperatures and longer durations in cold temperatures favor recovery. However, Purcell (1980) evaluated recovery of infected grapevines in a variety of locations selected to create an environmental cline and reported that winter recovery was not consistently related to lower winter temperatures.

In addition to identifying regional and cultivar differences in recovery rates to better understand existing Pierce’s disease epidemiology, associating specific parameters (e.g. number of degree days, specific pathologies) with likelihood of recovery is necessary to translate risk assessments to new disease introductions and to future climate conditions. Around the world, areas suitable for viticulture are expected to change, moving farther north and towards higher elevations as temperatures increase (Hannah et al. 2013). For existing grape-growing regions, increased temperatures impact phenology and fruit quality, as well as cultivar suitability in general (van Leeuwen et al. 2019). Conversations around adapting viticulture systems for climate change should incorporate disease considerations in addition to other management practices such as later-ripening grape varieties and irrigation (van Leeuwen et al. 2019). In addition, more research regarding the impact of abiotic stresses such as drought on winter recovery should be considered as water deficit can exacerbate Pierce’s disease symptoms (Choi et al. 2013).

Overall, this study highlights the need to define specific parameters used to quantify disease progression and plant recovery under different climate conditions (heat units, chill hours etc) so that broader comparisons can be made across separate research trials. The low rates of recovery observed also underscore the possibility that in many locations winter recovery from Pierce’s disease is likely to be minimal or non-existent. Under future climate scenarios, locations that currently experience some recovery may not, leading to increases in disease severity. Best practices in managing Pierce’s disease and other diseases caused by *X. fastidiosa* should therefore include some consideration of regional climate and predicted summer and winter temperature ranges.

## Acknowledgments

Funding for this work was from United States Department of Agriculture (USDA) Agricultural Research Service appropriated project 2034-22000-014-00D. Mention of trade names or commercial products in this publication is solely for the purpose of providing specific information and does not constitute endorsement by USDA. USDA is an equal opportunity provider and employer.

## Notes

### Competing Interest Statement

The authors have declared no competing interest.

